# Insight into the origin of 5’UTR and source of CpG reduction in SARS-CoV-2 genome

**DOI:** 10.1101/2020.10.23.351353

**Authors:** Ali Afrasiabi, Hamid Alinejad-Rokny, Nigel Lovell, Zhenming Xu, Diako Ebrahimi

## Abstract

SARS-CoV-2, the causative agent of COVID-19, has an RNA genome, which is, overall, closely related to the bat coronavirus sequence RaTG13. However, the ACE2-binding domain of this virus is more similar to a coronavirus isolated from a Guangdong pangolin. In addition to this unique feature, the genome of SARS-CoV-2 (and its closely related coronaviruses) has a low CpG content. This has been postulated to be the signature of an evolutionary pressure exerted by the host antiviral protein ZAP. Here, we analyzed the sequences of a wide range of viruses using both alignment-based and alignment free approaches to investigate the origin of SARS-CoV-2 genome. Our analyses revealed a high level of similarity between the 5’UTR of SARS-CoV-2 and that of the Guangdong pangolin coronavirus. This suggests bat and pangolin coronaviruses might have recombined at least twice (in the 5’UTR and ACE2 binding regions) to seed the formation of SARS-CoV-2. An alternative hypothesis is that the lineage preceding SARS-CoV-2 is a yet to be sampled bat coronavirus whose ACE2 binding domain and 5’UTR are distinct from other known bat coronaviruses. Additionally, we performed a detailed analysis of viral genome compositions as well as expression and RNA binding data of ZAP to show that the low CpG abundance in SARS-CoV-2 is not related to an evolutionary pressure from ZAP.

## Introduction

Severe Acute Respiratory Syndrome Coronavirus 2 (SARS-CoV-2), the causative agent of COVID-19 pandemic, has a single-stranded positive RNA (+ssRNA) genome of ~30kb long, which is one of the largest known viral RNA genomes (Romano, et al. 2020). SARS-CoV-2 RNA has unique genomic features with likely roles in the high pathogenicity and cross-species transmission of this virus (Coronaviridae Study Group of the International Committee on Taxonomy of 2020; Gussow, et al. 2020). An expansive view of these genomic features is essential to improve our current understanding of the source and evolutionary path of this virus.

So far, comparative genomic studies suggest that, overall, SARS-CoV-2 is closely related to RaTG13, which is a coronavirus isolated from a species of bat known as the *intermediate horseshoe bat* (*Rhinolophus affinis*), in the Yunnan Province of China (Jaimes, et al. 2020; Zhou, Chen, et al. 2020; Zhou, Yang, et al. 2020). However, the genomic region coding for the ACE2 cellular receptor binding domain of SARS-CoV-2 Spike protein is an exception in that it is highly similar to the sequence of a Sunda pangolin (*Manis javanica*) coronavirus found in the Guangdong Province (Frutos, et al. 2020; Lam, et al. 2020; Xiao, et al. 2020). Both the sequence divergence of SARS-CoV-2 genome and its unique features including a 12-base insert creating a polybasic cleavage site, have sparked extensive debates on the origin of this new coronavirus (Andersen, et al. 2020; Boni, et al. 2020; Frutos, et al. 2020; Lam, et al. 2020; Leitner and Kumar 2020; Xiao, et al. 2020). Earlier studies suggest that SARS-CoV-2 was originated in bat, and pangolin was the intermediate species for its leap into human (Lam, et al. 2020; Liu, et al. 2020; Wahba, et al. 2020; Xiao, et al. 2020; Zhang, et al. 2020), However, a recent study based on phylogenetic dating methods indicate that the origin of ACE2 binding domain in the SARS-CoV-2 genome is likely a bat coronavirus, which is yet to be sampled (Boni, et al. 2020).

One of the unique genomic features of SARS-CoV-2 is the low abundance of CpG (di Gioacchino, et al. 2020; Pollock, et al. 2020; Wang, et al. 2020; Xia 2020). CpG suppression is a well-known phenomenon in viruses particularly those with RNA genomes (Karlin, et al. 1994; Rima and McFerran 1997). It has been reported that the CpG composition of +ssRNA viral genomes often mimics the CpG content of their hosts (Cheng, et al. 2013). This suggests host CpG manipulating mechanisms play a role in shaping +ssRNA viral genomes during cross-species transmission. Nevertheless, these molecular mechanisms are not fully understood. One of the suggested mechanisms is DNA cytosine methylation-induced deamination (Bird 1980; Cheng, et al. 2013; Alinejad-Rokny, et al. 2016). However, SARS-CoV-2 does not have a DNA stage, thus this mechanism is not likely to be relevant. Another suggested mechanism is recognition of CpG sites within viral RNA by the host RNA-binding protein ZAP (CCCH-type zinc finger antiviral protein) (Xia 2020). ZAP is known to restrict viral replication by binding to the CpG rich regions of viral RNA, and subsequently inducing viral RNA degradation (Gao, et al. 2002; Ghimire, et al. 2018). The low CpG content of RNA viruses have been proposed to be linked to this ZAP-dependent depletion of CpG-rich viral RNA sequences (Gao, et al. 2002; Takata, et al. 2017; Ghimire, et al. 2018). Nevertheless, ZAP, despite having a broad antiviral role, does not restrict all viruses (Ghimire, et al. 2018).

A recent study by Xia suggested that the low CpG content of SARS-CoV-2 might be due to an evolutionary pressure from ZAP (Xia 2020). Among the coronaviruses studied in Xia’s study, those isolated from canines were shown to have the most CpG depleted genomes. Therefore, it was postulated that dogs may have been the intermediate species for the emergence of SARS-CoV-2. Furthermore, based on a previous report that ZAP identifies CpG-containing RNA (Takata, et al. 2017), Xia linked the low CpG content of SARS-CoV-2 genome to the high expression level of ZAP in the intermediate host tissues (Xia 2020). This hypothesis is based on two assumptions, which are not likely to be correct: first, only the frequency of CpG (but not those of other 15 dinucleotides ApA, ApC, …, TpT) is sufficient to make inferences about the origin of viruses, and second, ZAP is the main selection force against viral CpGs. A number of follow up studies have challenged Xia’s methodology and conclusion. For instance, Pollock *et al*. repeated Xia’s analysis using a larger number of SARS-CoV-2 related viruses and found that CpG deficiency is not specific to dog coronaviruses or SASR-CoV-2, and it is observed in pangolin coronaviruses and to a greater extent in pangolin pestiviruses (Pollock, et al. 2020). Moreover, modeling of the binding affinity of ACE2 and SARS-CoV-2 Spike protein in 410 vertebrates showed a low score of susceptibility to SARS-CoV-2 infection for dogs (Damas, et al. 2020). This finding was also confirmed by viral replication experiments (Shi, et al. 2020). Digard *et al*. showed that CpG abundance varies significantly across the SARS-CoV-2 genome, with envelop and ORF10 not showing CpG depletion. They showed that the CpG levels of SARS and SARS-CoV-2 envelop sequences are even higher than those of envelop from other human coronaviruses. Using a phylogenetic analysis, the authors argued that these genomic composition changes are more likely to be an ancestrally-driven traits related to the origin of these viruses in bats, not due to a post-zoonotic transfer selection force. These data suggest that the overall CpG content alone is not a reliable index for inferring the host origin of viruses (Digard, et al. 2020).

Here, we investigate the origin of SARS-CoV-2 using a comparative genomic approach. Additionally, we perform a detailed analysis of multiple data sets including the representations of short sequence motifs in viral genomes, ZAP expression, and CpG motif preference of ZAP to investigate the role of ZAP in reducing SARS-CoV-2 CpG level. Our analyses show that the 5’UTRs of SARS-CoV-2 and Guangdong pangolin coronavirus (accession number EPI_ISL_410721) are highly similar. This suggests bat and pangolin coronaviruses might have recombined at least twice (in the 5’UTR and ACE2 binding regions) to seed the formation of SARS-CoV-2. An alternative hypothesis is that the lineage preceding SARS-CoV-2 is a yet to be sampled bat coronavirus whose ACE2 binding domain and 5’UTR are distinct from other known bat coronaviruses. We show that the representation of almost all dinucleotides, not only CpG is different in SARS-CoV-2 compared to its more distantly related coronaviruses. For example, GpC, TpC, and ApT are all represented at significantly lower levels in the SARS-CoV-2 genome compared to the SARS-CoV genomes. Our analyses indicate that the CpG motifs preferentially targeted by ZAP do not have a lower representation than those not often recognized by ZAP. Altogether, our results provide multiple lines of evidence against the role of ZAP in shaping the SARS-CoV-2 genome.

## Results

### Analysis of motif representations using PCA

We applied principal component analysis (PCA) on motif representation (D-value) matrices to interrogate similarities between and within virus families (**Supplementary Table 1 and 2**). **Fig. 1A**shows the PCA of all four virus groups studied here, and **Fig. 1B**shows the results of the same analysis done on coronaviruses only. As indicated in **Fig. 1A**, all of the four virus families HIN1, HBV, HIV-1 and coronaviruses are separated using the first two PCs. PCA of only coronavirus sequences is depicted in **Fig. 1B**. As expected, SARS-COV-2, Bat-RaTG13, and Pangolin-CoV formed a cluster (SARS-CoV-2-like group), which was separated from the rest of coronavirus sequences, Human-SARS-CoV, Paguma-SARS-CoV, Viverridae-SARS-CoV, Paradoxurus-SARS-CoV, Bat-SARS-CoV, Mus-SARS-CoV and Primate-SARS-CoV (SARS-CoV group). The two groups of coronaviruses described above (shown in ovals in **Fig. 1B**) are clearly separated from each other, except for three bat coronaviruses shown by black arrows (accession numbers MG772933, MG772934 and KY352407). In the PC1-PC2 plot of **Fig. 1B** these three sequences are positioned closer to the SARS-CoV-2-like group than to their own SARS-CoV group. Alignment-based comparative genomic studies have shown that SARS-CoV-2 sequences are most closely related to the Bat-RaTG13 sequence (Zhou, Chen, et al. 2020). This is also apparent in the PC1-PC2 plot of **Fig. 1B** with Bat-RaTG13 (blue dot) being the closest to the SARS-CoV-2 cluster (red dots). Pangolin-CoV sequences (green dots) also position close to the SARS-CoV-2 sequences (**Fig. 1B**). The Pangolin-CoV sequence with the accession number EPI_ISL_410721, which is labeled in PubMed to be of Guangdong origin, is the second most closely related virus to the SARS-CoV-2 family. The remaining Pangolin-CoV sequences, which are labeled with Guangxi, form a sub-cluster that is located at a farther distance to the SARS-CoV-2 sequences.

**Figure 1:**
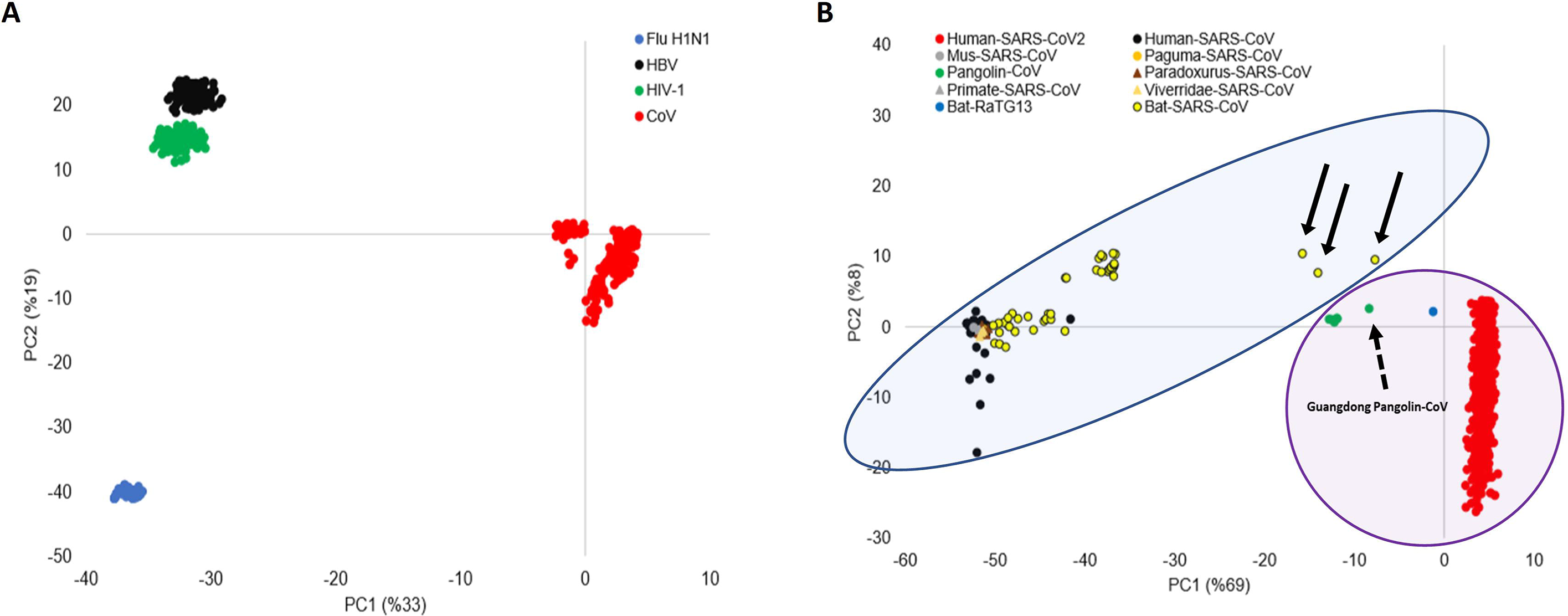
PCA of viral motifs representations. D-values of all dinucleotide and trinucleotide motifs in all viral sequences form a matrix, which is used as an input for PCA. (A) PC1-PC2 plot shows four clusters, one for each virus family: H1N1, SARS-CoV, HBV, and HIV-1. (B) PC1-PC2 plot classifies SARS-CoV viruses into two clusters. SARS-COV-2, Bat-RaTG13, and Pangolin-CoV formed a cluster (SARS-CoV-2-like group), which is separated from the rest of coronavirus sequences Human-SARS-CoV, Paguma-SARS-CoV, Viverridae-SARS-CoV, Paradoxurus-SARS-CoV, Bat-SARS-CoV, Mus-SARS-CoV and Primate-SARS-CoV (SARS-CoV group). SARS-CoV-2-like and SARS-CoV groups are highlighted with purple and blue circles. The three Bat-SARS-CoV sequences are located close to the SARS-CoV-2-like group, which is shown by black arrows. Guangdong Pangolin-CoV is shown by a broken arrow.

### 5’UTR sequence similarity

To investigate the origin of SAR-CoV-2, we compared, using a moving window analysis, its genome to the sequences of Bat-RaTG13 and two Pangolin-CoV genomes, one from Guangdong (accession number EPI_ISL_410721), and one from Guangxi (accession number EPI_ISL_410539). The results are shown in **Fig. 2**, which depicts the difference between pairwise similarities of SARS-CoV-2/Bat-RaTG13 and SARS-CoV-2/Pangolin-CoV in windows of size 20bp moving along the sequence by 1bp. Windows within which Bat-RaTG13 sequence is more closely similar to SARS-CoV-2 than Pangolin-CoV to SARS-CoV-2 are shown in blue vertical bars, and the opposite is shown in red. As expected, the majority of windows are blue indicating that, overall, Bat-RaTG13 is the most similar virus to SARS-CoV-2. The windows indicated in red are those within which Pangolin-CoV is more similar to SARS-CoV-2. Overall, for both Guangdong and Guangxi sequences there are regions within which these sequences are more similar to SARS-CoV-2 (6% and 9% of windows, red bars in **Fig. 2** for Pangolin Guangxi and Guangdong, respectively). However, there are many more windows within which SARS-CoV-2 is more similar to Bat-RaTG13 (94% and 91% of windows, blue bars in Pangolin Guangxi and Guangdong, respectively panels in **Fig. 2**). The ACE2-binding domain of SARS-CoV-2 Spike is known to have a very high sequence similarity to the same region in the Guangdong pangolin coronavirus genome (Frutos, et al. 2020; Lam, et al. 2020; Xiao, et al. 2020). This is also indicated by two sharp red peaks in **Fig. 2B** and to a lesser extent in **Fig. 2A**. The nucleic acid as well as amino acid sequences of these two regions are shown in boxes above the similarity lines. Most importantly, our window analysis shows a region within 5’UTR that is identical in SARS-CoV-2 and Guangdong pangolin coronavirus sequences but differs in five positions between Bat-RaTG13 and SARS-CoV-2 sequences. These genomic positions are 19, 37, 91, 174 and 190, relative to the SARS-CoV-2 reference sequence (accession number NC_045512). Such a high level of similarity between the Guangdong pangolin coronavirus and SARS-CoV-2 is not observed for the Guangxi pangolin coronavirus. To investigate if the high level of similarity between the 5’UTRs of SARS-CoV-2 and Guangdong pangolin coronavirus is also observed for other coronaviruses, we performed a multiple alignment of 505 sequences of coronaviruses from a wide range of hosts such as human, pangolin, bat, bovine, canine, camel, and rabbit. A section of this alignment, shown in **Supplementary Fig 1**, indicates that Guangdong pangolin coronavirus and SAR-CoV-2 have the most similar 5’UTRs among all reported coronaviruses.

**Figure 2:**
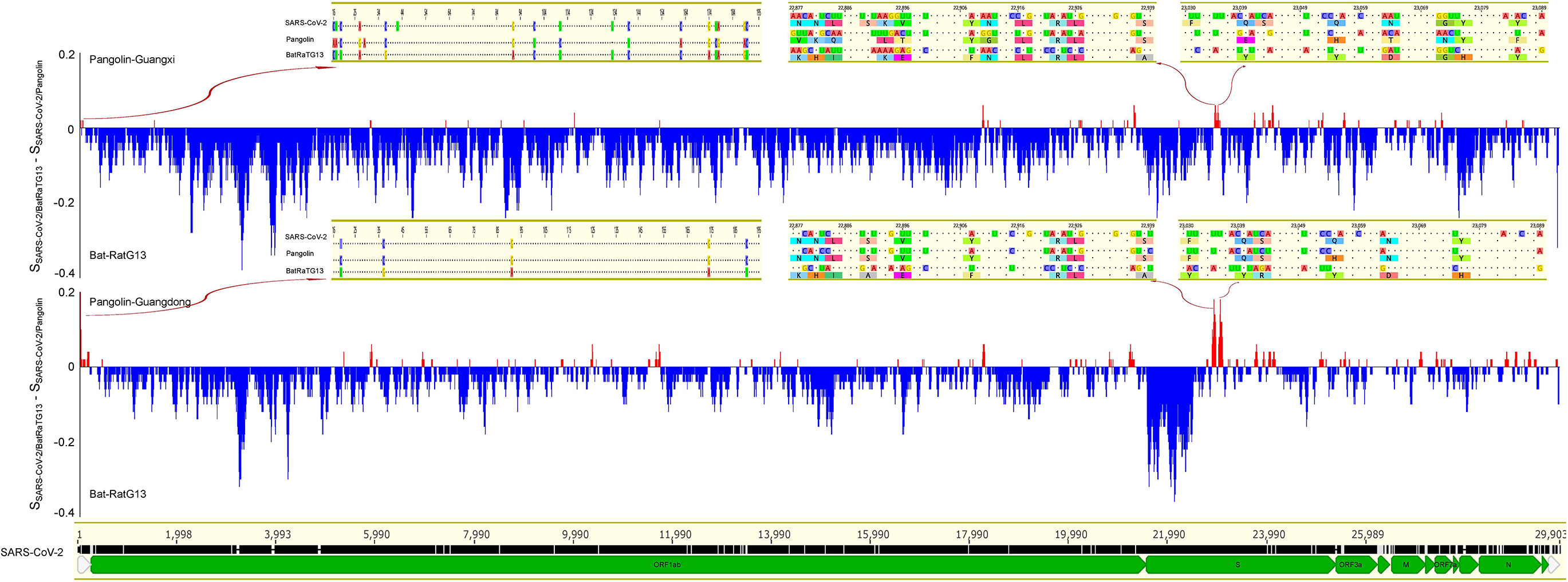
5’UTR sequence similarity between SARS-CoV-2 and pangolin-CoV. The sequences of Bat-RaTG13 and two Pangolin-CoV genomes, one from Guangdong (accession number EPI_ISL_410721), and one from Guangxi (accession number EPI_ISL_410539) were compared with the SARS-CoV-2 reference sequence. A 20bp sliding window moving by 1bp was used to quantify the difference between two pairwise similarity measures (SARS-CoV-2/Bat-RaTG13 and SARS-CoV-2/Pangolin-CoV). Windows within which Bat-RaTG13 sequence is more closely similar to SARS-CoV-2 than Pangolin-CoV is to SARS-CoV-2 are shown in blue vertical bars, and the opposite is shown in red.

As indicated in **Fig. 1**, coronaviruses used in this study form two distinct groups (SARS-CoV-2-like and SARS-CoV) based on their motif representations (D-values). We compared, for each dinucleotide, the median D-values of these two groups (**Fig. 3** and **Supplementary Table 3**). All dinucleotide motifs (not only CpG) except for ApG were significantly different between the two groups. There is an excess of ApA, TpA, GpT, TpT, ApC, CpC and GpG, and a deficit of CpA, GpA, ApT, CpT, TpC, GpC, TpG and CpG in group one (SARS-CoV-2-like) compared to group two (SARS-CoV). We then performed an additional representation analysis focusing on the trinucleotide motifs that contain CpG (**Fig. 4** and **Supplementary Table 3**). Our analysis shows that the trinucleotide profiles of the two virus groups are also different. Specifically, there is a higher representation for CGC, CGG, CCG, and ACG and lower representation for CGA, CGT, TCG and GCG in group one compared to group two coronaviruses.

**Figure 3:**
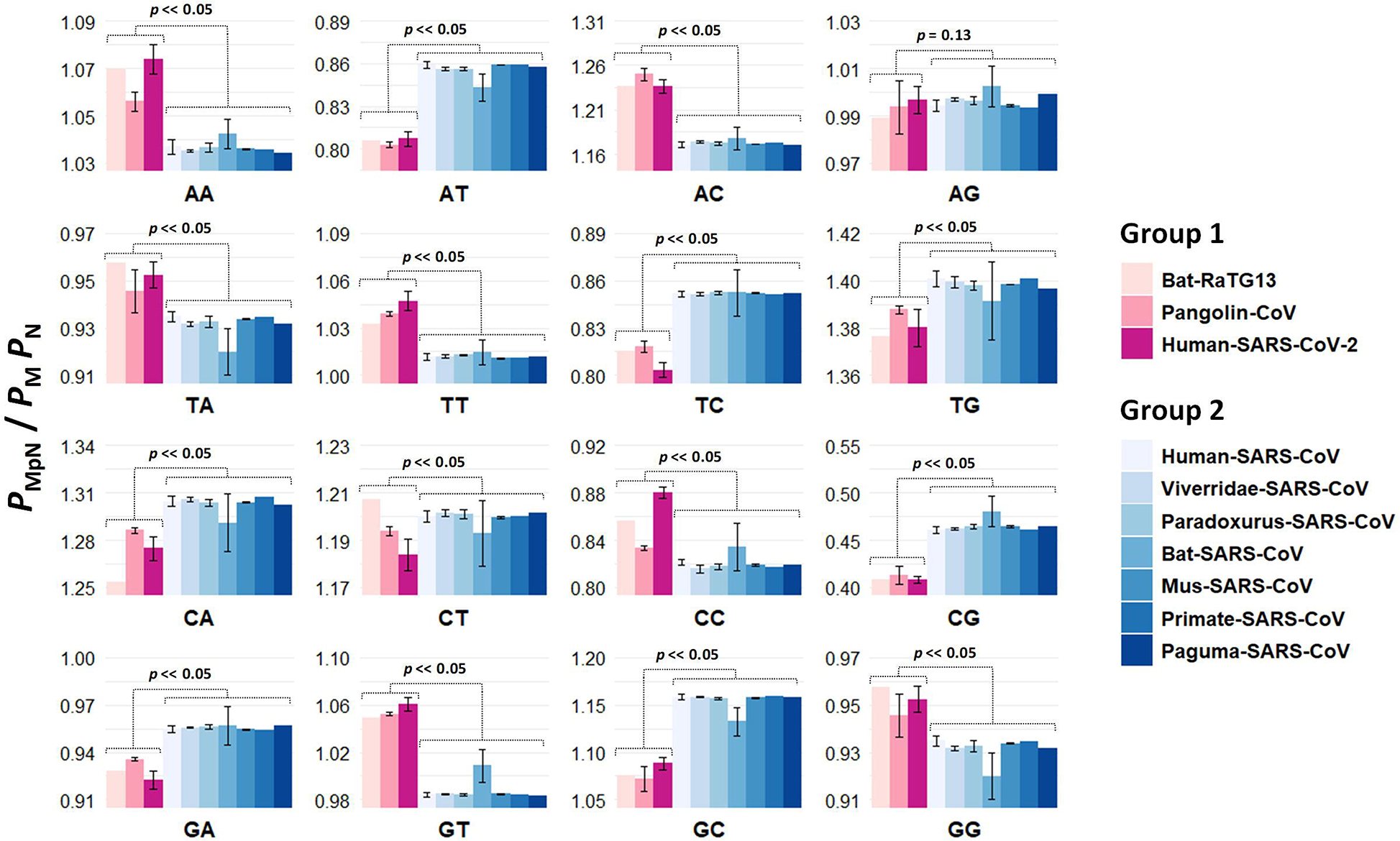
Comparison of dinucleotide motif representations between SRAS-CoV-2-like and SARS-CoV groups. D-values of each dinucleotide were compared between the two virus groups SARS-CoV-2-like (SARS-COV-2, Bat-RaTG13, and Pangolin-CoV) and SARS-CoV (Human-SARS-CoV, Paguma-SARS-CoV, Viverridae-SARS-CoV, Paradoxurus-SARS-CoV, Bat-SARS-CoV, Mus-SARS-CoV and Primate-SARS-CoV). Mann–Whitney test was used to examine the difference in the median of D-values between the two coronavirus groups.

**Figure 4:**
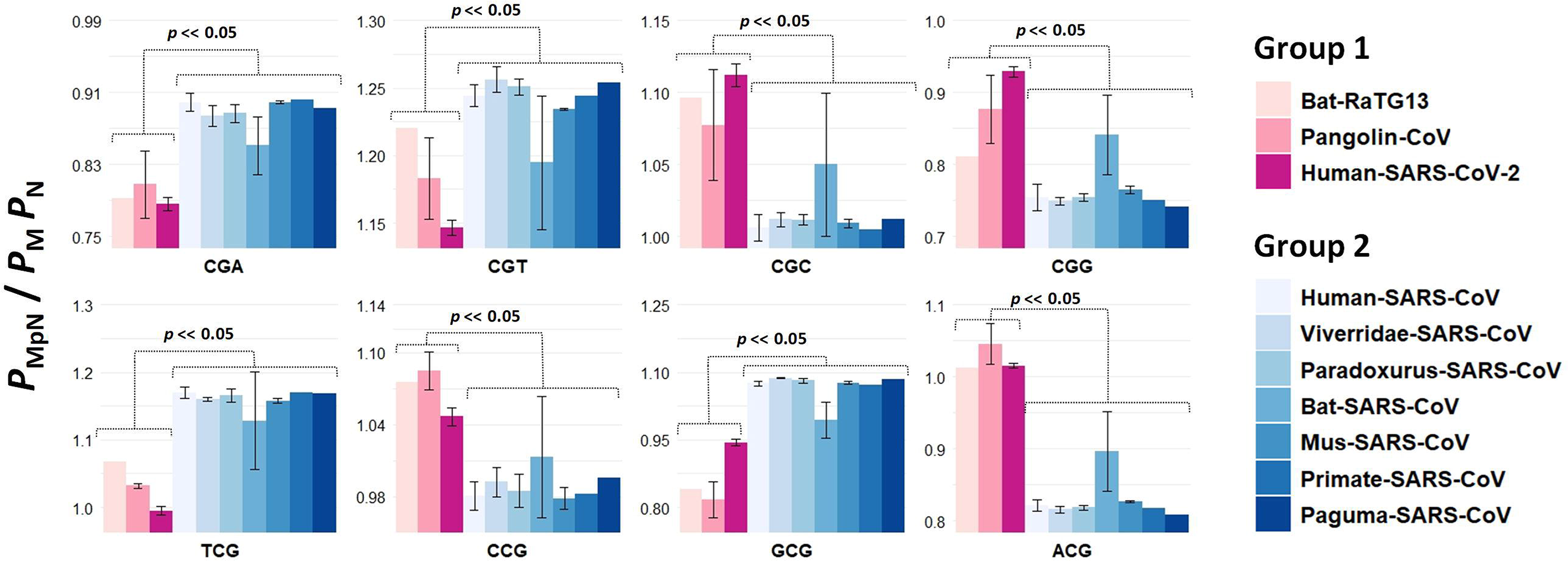
Comparison of CpG trinucleotide motif representations between SRAS-CoV-2-like and SARS-CoV groups. D-values of each trinucleotide were compared between the two virus groups SARS-CoV-2-like (SARS-COV-2, Bat-RaTG13, and Pangolin-CoV) and SARS-CoV (Human-SARS-CoV, Paguma-SARS-CoV, Viverridae-SARS-CoV, Paradoxurus-SARS-CoV, Bat-SARS-CoV, Mus-SARS-CoV and Primate-SARS-CoV). Mann–Whitney test was used to examine the difference in the median of D-value between the two coronavirus groups.

To determine if host ZAP level has played a role in altering viral CpG levels, we investigated the correlation between ZAP mRNA expression level and CpG motif representation in viral genomes (**Fig. 5** and **Supplementary Table 4**). To ensure our analyses are physiologically relevant, we used the ZAP expression data of common cellular/tissue reservoirs for each virus. We observed no significant correlation between CpG motif representation and ZAP mRNA expression (R^2^ = 0.04, *p* = 0.47, **Fig. 5**).

**Figure 5:**
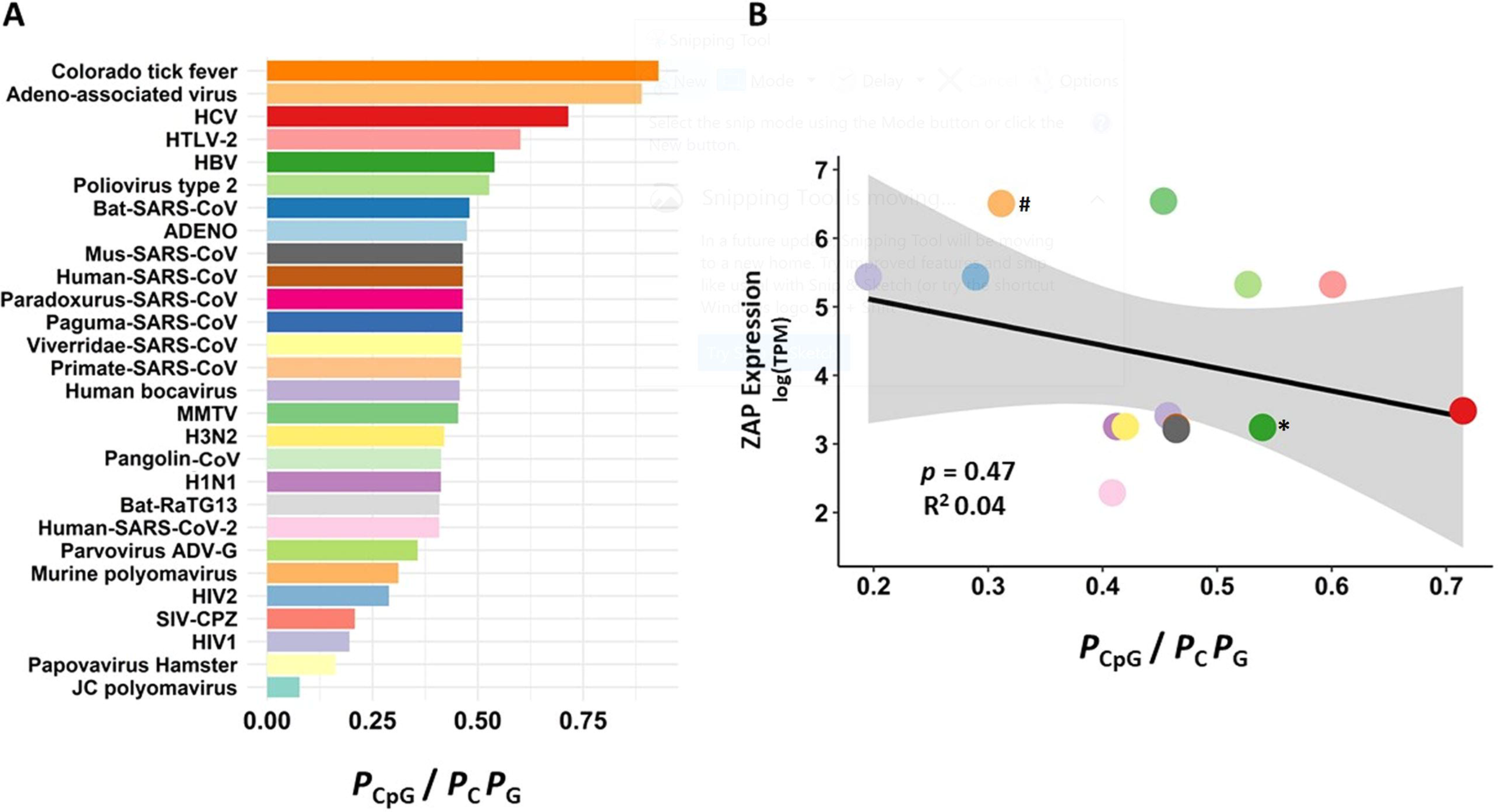
Correlation between ZAP mRNA level and viral CpG motif representation. (A) D-values of CpG motifs in different viruses. (B) Linear regression model was used to test the correlation between the CpG D-values of viral genomes and ZAP mRNA expression levels in common virus reservoirs. Colors match in panels A and B. All viruses in panel B are RNA viruses except HBV (*), which has a partially double stranded circular DNA genome and Murine polyomavirus (#), which is a double stranded DNA virus.

To determine the CpG motif specificity of ZAP, we overlaid the ZAP CLIP-seq data and the CpG distribution of two viruses, HIV and JEV (Japanese Encephalitis Virus) (**Fig. 6**). We found no clear and consistent pattern of co-localization between ZAP binding regions and CpG sites in HIV and JEV genomes.

**Figure 6:**
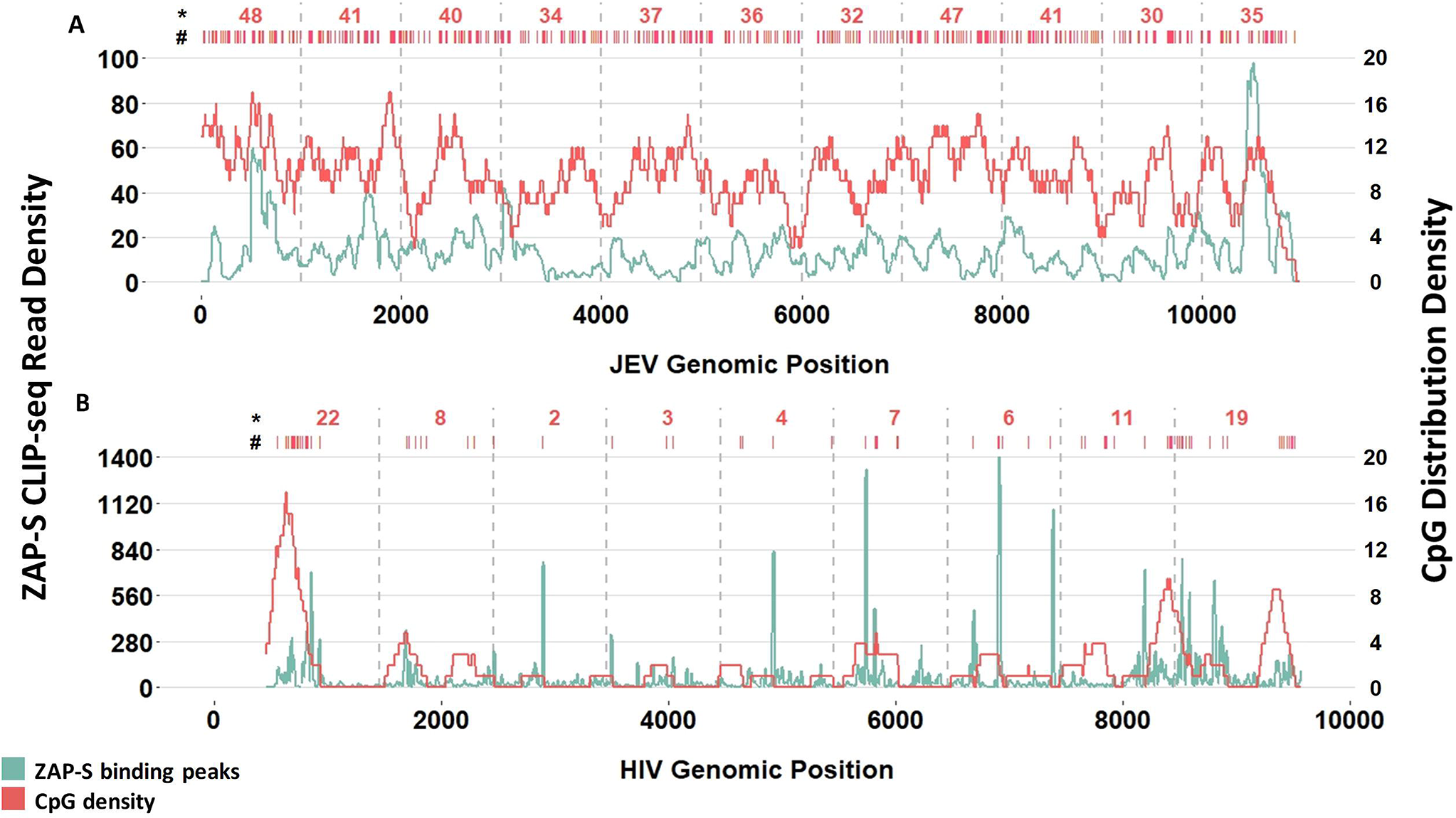
Co-location of ZAP binding regions and CpG motifs. Overlaying of the ZAP binding peaks and CpG densities in (A) JEV genome (Japanese Encephalitis Virus) and (B) HIV-1. The ZAP binding peaks (density of reads aligned to the genome) are estimated using a 250bp sliding window moving by 1bp along the viral genomes. The CpG density was calculated using the same sliding window analysis method, except we used a 200bp window sliding by 1bp in JEV and a 250bp window sliding by 1pb in HIV-1. ZAP binding peaks and CpG densities are shown in green and red, respectively. # Location of CpGs. * Number of CpGs per 1Kb.

It has been shown that ZAP binds preferentially to C(n7)G(n)CG motifs (Luo, et al. 2020). To further investigate the role of ZAP binding in reducing SARS-CoV-2 CpG level, we compared the relative frequency of C(n7)G(n)CG in group one (SARS-CoV-2-like) and group two (SARS-CoV) viruses (**Supplementary Table 5**). In spite of CpG reduction in SARS-CoV-2 genome, the ZAP preferred motif C(n7)G(n)CG, is more abundant in the SARS-CoV-2-like group (**Fig. 7**).

**Figure 7:**
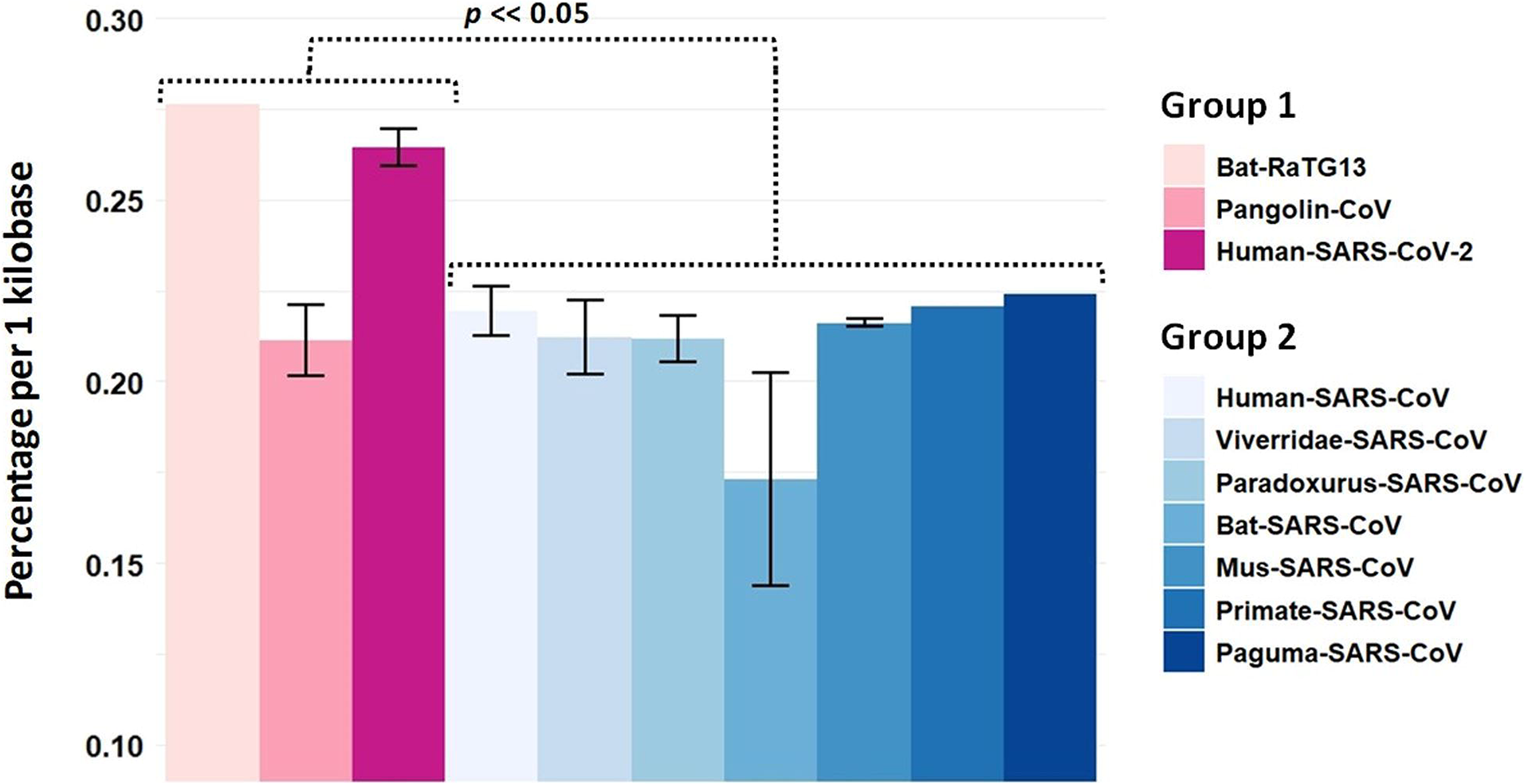
Comparison of the abundance of ZAP optimal binding motif C(n7)G(n)CG in viruses of SARS-CoV-2-like and SARS-CoV groups. The percentages of ZAP optimal binding motif C(n7)G(n)CG among all CpG motifs per 1kb in the two coronavirus groups we compared. The SARS-CoV-2-like group includes SARS-COV-2, Bat-RaTG13, and Pangolin-CoV. The SARS-CoV group includes Human-SARS-CoV, Paguma-SARS-CoV, Viverridae-SARS-CoV, Paradoxurus-SARS-CoV, Bat-SARS-CoV, Mus-SARS-CoV, and Primate-SARS-CoV. Mann–Whitney test was used to determine the difference in the median of D-values between these two coronavirus groups.

In Xia’s study, it is shown that the genome of one of the dog coronaviruses (accession number KC175339) has a significantly higher CpG level than the rest of canine coronaviruses. The author attributed the high CpG level of this virus to its extensive propagation in cell culture before it is sequenced. Xia argues that lack of a selection force against CpG has led to its rebound in this virus (Xia 2020). To test this rebounding hypothesis, we quantified the CpG motif representations of HIV-1 lab strain IIIB (accession numbers A04321.1), which is commonly cloned and cultured in cell lines, and compared it with the CpG representation of HIV sequences isolated from patients. **Fig. 8** indicates the distribution of CpG representations in 844 HIV-1 isolate sequences. The CpG representation values for A04321.1 falls within the interquartile range, thus is not significantly different from the median of this distribution.

**Figure 8:**
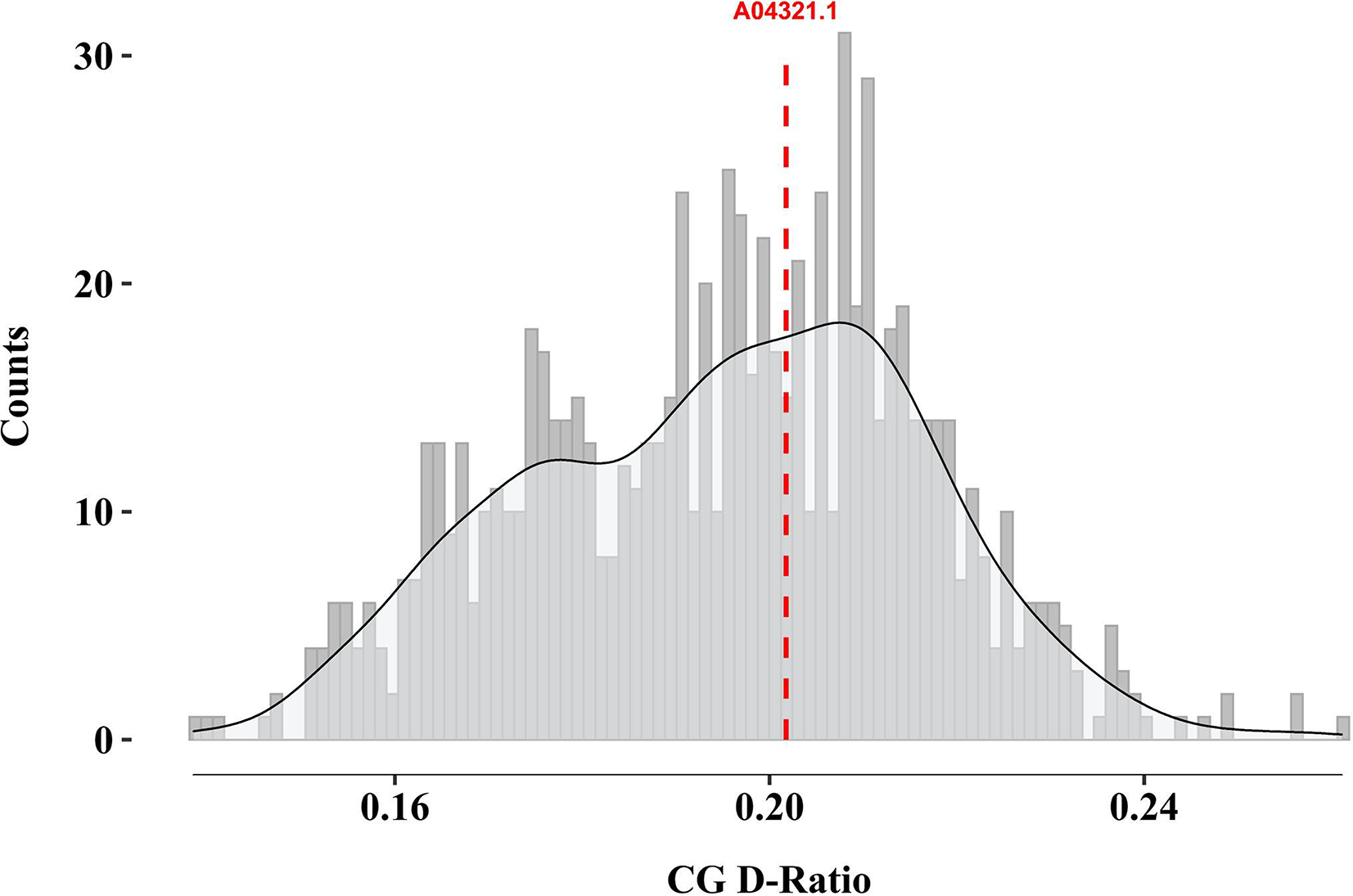
Comparison of CpG abundance in the HIV-1 molecular clone IIIB and HIV-1 isolates obtained from patients. A total of 844 HIV-1 sequences were used to generate a distribution of CpG D-values. The CpG D-value of HIV-1 IIIB (accession number A04321.1) is shown by a vertical dashed line.

## Discussion

It is critical to determine the origin of SARS-CoV-2 and understand the changes leading to its leap into human. This information can help prevent and/or better control future pandemics caused by coronaviruses. Alignment-based comparative genomic studies indicate that the shoehorse bat coronavirus Bat-RaTG13 is the most closely related virus to SARS-CoV-2. However, the ACE2 biding domain sequence of SARS-CoV-2 is more similar to that of a Guangdong pangolin coronavirus. These results led to the conclusion, in several independent studies, that SARS-CoV-2 originated via a recombination between the bat and pangolin coronaviruses (Lam, et al. 2020; Li, et al. 2020; Xiao, et al. 2020). However, a recent study (Boni, et al. 2020) shows that phylogenetic dating methods do not support the bat-pangolin recombination hypothesis. The study suggests that the origin of SARS-CoV-2 is likely a coronavirus circulating in bats, but this virus remains to be sampled.

We have previously shown that an alignment-free method that uses the representation of short sequence motifs can precisely identify HIV-1 subtypes (Ebrahimi, et al. 2014). Here, we used the same method to investigate the similarities between and within virus families. As indicated in **Fig. 1A**, this method successfully classifies different virus families. Additionally, it shows a separate cluster for SARS-CoV-2 with Bat-RaTG13 and pangolin coronaviruses in its close proximity (see SARS-CoV-2-like group in **Fig 1B**). Our method confirmed that SARS-CoV-2 sequences are closely similar to Bat-RaTG13 (MN996532) followed by a Guangdong pangolin coronavirus (EPI_ISL_410721). Additionally, our window analysis revealed that the 5’UTR of SARS-CoV-2 is highly similar to the 5’UTR of Guangdong pangolin coronavirus. Altogether, our analyses indicate that both the 5’UTR and ACE2-binding domain of SARS-CoV-2 genome have either a pangolin coronavirus origin (Lam, et al. 2020; Xiao, et al. 2020) or they have originated from an unsampled bat coronavirus, which is more closely related to SARS-CoV-2 than both RaTG13 and Guangdong pangolin coronavirus are to SARS-CoV-2 (Boni, et al. 2020). Regardless of which hypothesis is correct, and what coronavirus (bat or bat-pangolin recombinant) was the immediate predecessor of SARS-CoV-2, our results suggest that recombinations of both 5’UTR and ACE2 binding regions of coronaviruses are relatively recent events in the history of SARS-CoV-2.

Our analyses confirmed that SARS-CoV-2 and its closely related viruses Bat-RaTG13 and Pangolin-CoV have a lower CpG content compared to other viruses (**Fig. 3**) (di Gioacchino, et al. 2020; Pollock, et al. 2020; Wang, et al. 2020; Xia 2020). However, we show that changes in motif representation is not exclusive to CpG. All dinucleotides (except for ApG) have significantly different representations in the viruses of SARS-CoV-2-like group compared to the viruses in the SARS-CoV group (**Fig. 3**). Not only CpG, but also CpA, GpA, ApT, CpT, and TpC have lower representations in SARS-CoV-2 and its closely related viruses. Additionally, ApA, TpA, TpT, GpT, ApC, CpC, and GpG have higher representations in the viruses of SARS-CoV-2-like group. The tri-nucleotide profiles of these two virus groups are also totally distinct (**Fig. 4**). Altogether, these data suggest that the observed differences in CpG abundance between the two virus groups is part of a global genomic difference rather than being a signature of an exclusive selection force against CpG motifs.

Immune evasion is one of the mechanisms proposed to explain the low CpG abundance in viral genomes including coronavirus sequences (Shpaer and Mullins 1990; Karlin, et al. 1994; Shackelton, et al. 2006; Woo, et al. 2007; Cheng, et al. 2013; Greenbaum, et al. 2014). Assuming that this hypothesis is true, one would expect to observe a significant CpG depletion in the genome of SARS-CoV-2 to justify its high transmission rate and pathogenicity. By contrast, there is a little difference between the SARS-CoV-2-like and SARS-CoV viruses in terms of CpG abundance. The average CpG counts per kilobase (Kb) is 14.7 and 19.3 in group one (SARS-CoV-2-like) and group two (SARS-CoV) viruses, respectively. This means, on average, compared to SARS-CoV, SARS-CoV-2 has ~4.6 less CpGs per 1 Kb. It is unlikely that such a marginal CpG difference plays a critical role in SARS-CoV-2 immune evasion. Importantly, a previous study showed no correlation between the pathogenicity and global CpG content of coronaviruses infecting humans (di Gioacchino, et al. 2020). Furthermore, it has been shown that the CpG content of SARS-CoV-2 sequences is highly variable across the viral genome. In some regions of the SARS-CoV-2 genome such as envelop, not only CpG is not depleted, but it is more abundant compared to some of the other coronaviruses. This suggests the global CpG content is unlikely to be a vital genomic feature of SARS-CoV-2 with a possible role in immune evasion (Digard, et al. 2020). Altogether, the overall reduction of CpG level in the SARS-CoV-2 genome is likely unrelated to the pressure imposed on the virus by the innate immune system.

Among the components of human innate immune system, ZAP has been shown to play a key role in the inhibition of RNA viruses by binding to CpG containing sequences and recruiting a RNA-degradation machinery (Takata, et al. 2017). A recent study shows that ZAP is capable of inhibiting SARS-CoV-2 *in vitro* (Nchioua, et al. 2020). Xia’s study postulates that the source of SARS-CoV-2 is a bat coronavirus whose genome underwent further CpG reduction by ZAP after the virus infected an intermediate species with a high ZAP expression level (possibly a canine tissue). It was suggested that ZAP-induced CpG depletion of viral RNA in this intermediate species led to the generation of SARS-CoV-2, which was able to infect human cells (Xia 2020). We tested this hypothesis using the CpG representation data of multiple viruses and the expression data of ZAP in tissues infected by these viruses. As indicated in **Fig. 5** we observe no correlation between the abundance of CpG in viral genomes and the ZAP mRNA level in viral reservoirs.

Further, our analyses show a poor association between the abundance of CpG across viral genomes and the location of ZAP binding peaks (**Fig. 6**). These data suggest that binding of ZAP to viral RNA is likely not governed by the global CpG abundance of viral genomes. In support of our results, previous studies have shown that ZAP does not inhibit all viruses and its inhibition is independent of the viral CpG content (Bick, et al. 2003). Additionally, a recent study shows that the location and sequence context of CpGs but not the overall CpG abundance of viral genome play a role in inducing ZAP antiviral activity (Ficarelli, et al. 2020). Moreover, one of the mechanisms by which ZAP inhibits viruses is through the suppression of viral mRNA translation via blocking eIF4A (Ghimire, et al. 2018). This mechanism is independent of ZAP binding to viral RNA.

A previous study has shown that mouse ZAP preferentially binds to C(n7)G(n)CG motifs where n: A, C, G, or U (Luo, et al. 2020). Assuming that human ZAP has the same motif preference and that it has induced an evolutionary pressure on the SARS-CoV-2 genome, one would expect the relative abundance of C(n7)G(n)CG to be lower in SARS-CoV-2 compared to other coronaviruses. Our analysis does not show a lower abundance for this motif in the SARS-CoV-2 genome (**Fig. 7**). In fact SARS-CoV-2 genome has a slightly higher %C(n7)G(n)CG. This may be considered as yet another line of evidence against the role of ZAP in lowering the CpG content of SARS-CoV-2 genome.

Xia’s study postulated that in the absence of ZAP, pressure against CpG is relieved and viruses evolve to restore their lost CpGs (Xia 2020). We tested this hypothesis using the HIV-1 molecular clone IIIB. Our analysis did not show a significant difference between the CpG content of HIV IIIB and those of HIV sequences isolated from patients (**Fig. 8**). This suggests the abundance of CpG motif in viral genomes is likely independent of ZAP-induced immune response in the virus reservoir.

We note that some of the studies of the evolutionary footprint of host immune mechanisms on viral genomes, focus merely on specific motifs, and ignore the overall composition of viral genomes. In many cases, this can lead to gross misinterpretation of data. For example, a phylogeny effect can be misinterpreted as an evolutionary signature. To better understand the role of ZAP and other restriction factors in the inhibition and/or evolution of viruses, a global analysis of viral genomic composition is needed. Differences observed in 15 out of 16 dinucleotides (i.e. not only CpG) (**Fig. 3**) point to general mechanism(s) with a global impact on the overall composition of SARS-CoV-2 genome. One of these mechanisms might be oxidative stress, although there is currently no data to support this hypothesis. Viral infection is often associated oxidative stress, which results in producing reactive oxygen species (ROS) (Schwarz 1996). It has been shown that coronavirus infection causes a high level of ROS production in host cells (Delgado-Roche and Mesta 2020). Nucleotides, particularly guanine, are more prone to the oxidative damage caused by ROS, which oxidize guanine to 8-oxyguanine in both DNA and RNA (Schneider, et al. 1993). 8-oxyguanine has a similar affinity for binding to adenine and cytosine (Sekiguchi and Tsuzuki 2002). There is a possibility that during the SARS-CoV-2 replication process (Romano, et al. 2020), which includes synthesizing a negative strand from the genome followed by making a positive strand using the newly synthesized negative strand, G is substituted with T. The lower representations of CpG, TpG, and GpA accompanied with higher representations of CpT, TpT and TpA in the viruses of SARS-CoV-2-like group (**Fig. 3**) might be due to G>T mutations induced by oxidative stress. Nevertheless, there is currently no data to support this hypothesis.

In summary, we performed several independent analyses to investigate similarities between SARS-CoV-2 and other coronaviruses and also to determine if ZAP played a role in the emergence of this virus. We discovered a high degree of similarity between the 5’UTRs of SARS-CoV-2 and Guangdong pangolin coronavirus. It remains to be determined if the high pathogenicity and transmissibility of SARS-CoV-2 is due, at least in part, to its unique 5’UTR sequence. Our analyses find no evidence to suggest that ZAP exerts an evolutionary pressure on the SARS-CoV-2 genome by targeting its CpG motifs. This, however, does not imply ZAP plays no role in inhibiting SARS-CoV-2.

## Methods

### Viral sequences acquisition

We analyzed a total of 3,967 full-length genomic sequences from different coronaviruses: 3,621 Human-SARS-CoV, 1 Paguma-SARS-CoV, 3 Viverridae-SARS-CoV, 6 Paradoxurus-SARS-CoV, 45 Bat-SARS-CoV, 41 Mus-SARS-CoV, 1 Primate-SARS-CoV, 242 Human-SARS-CoV-2, 1 Bat-RaTG13, 6 Pangolin-CoV. In addition, full-length sequences of 2,021 HIV-1, 12 HIV-2, 11 Adeno-associated virus, 4 Adeno-associated virus-1, 11 Colorado tick fever virus, 91 Flu H1N1 virus, 7365 H3N2 virus, 141 HBV, 6 HCV, 8 HTL-V2, 6 Human bocavirus, 576 JC polyomavirus, 3 Mouse mammary tumor virus, 9 Murine polyomavirus, 2 Papovavirus Hamster, 2 Parvovirus ADV-G, 4 Poliovirus type 2, and 7 SIV-CPZ were analyzed. GenBank (www.ncbi.nlm.nih.gov) was used to retrieve all 14,246 viral sequences (**Supplementary Table 1**).

### Analysis of motif representation

We quantified the representation of short sequence motifs (di- and tri-nucleotides) in all viral genomes using our previously reported D-value method (Ebrahimi, et al. 2011, 2012). Briefly, motif representation (D-value) is defined as the ratio of the observed frequency (*P*__obs__) of a motif over its expected frequency (*P*__exp__). *P*__obs__ is simply the observed relative frequency of the motif. *P*__exp__ is the total count of the motif in the sequence divided by the total count of all other motifs with the same length. An example of a D-value for a tri-nucleotide sequence is given in Eq. 1.

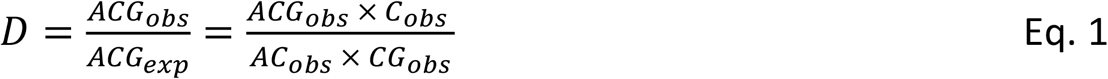

To demonstrate the ability of our nonalignment-based D-value method in classifying viral families we performed principal component analysis (PCA). D-values of all dinucleotide and trinucleotide motifs in all sequences form a matrix, which is used as an input for PCA.

We performed two separate principal component analyses. The first analysis was performed on all coronaviruses as well as flu H1N1, HIV, and HBV sequences to determine if analysis of motif representations can distinguish these viruses from coronaviruses. The second PCA was performed on coronavirus sequences to identify viruses whose genome show high similarities to the SARS-CoV-2 genome. Mann–Whitney test was used to determine the difference in the median of D-value between these two groups of coronaviruses.

### Analysis of association between CpG abundance and ZAP level

We interrogated the association between viral CpG representation (D-value) and *ZC3HAV1* (ZAP) mRNA expression level using a linear regression model. For viruses with more than one sequence we used an average D-value in our analyses. *ZC3HAV1* mRNA expression level data of common reservoirs for Human-SARS-CoV-2, Human-SARS-CoV, Mus-SARS-CoV, Paguma-SARS-CoV, HIV1, HIV2, H1N1, H3N2, HCV, HBV, Human bocavirus, Murine polyomavirus, Poliovirus type 2 and MMTV viruses were obtained from FANTOM5 (Lizio, et al. 2015).

To investigate the association of ZAP binding regions in the viral genomes with the number of CpGs co-located with these binding regions, we used the publicly available datasets of cross-linking immunoprecipitation (CLIP-seq) reported for ZAP. We obtained the processed ZAP CLIP-seq data (density of reads aligned to the genome) for wild type JEV (Japanese Encephalitis Virus) and wild type HIV (Takata, et al. 2017; Chiu, et al. 2018). We then calculated the CpG density across the JEV and HIV genomes using a sliding window analysis method. We used 200bp window and 1Bp sliding for JEV and 250bp window and 1Bp sliding for HIV.

The motif C(n7)G(n)CG has recently been demonstrated to be the optimal binding motif for Mouse ZAP (Luo, et al. 2020). To further investigate the role of ZAP in reducing CpG abundance in shaping the genome of SARS-CoV-2, we compared the relative abundance of C(n7)G(n)CG (#C(n7)G(n)CG / #CG, reported per kilobase) in the two viral groups of SARS-CoV-2-like and SARS-CoV. We used Mann–Whitney test to determine the difference between the two coronavirus groups).

## Supplementary Material

**Supplementary Fig. 1.**
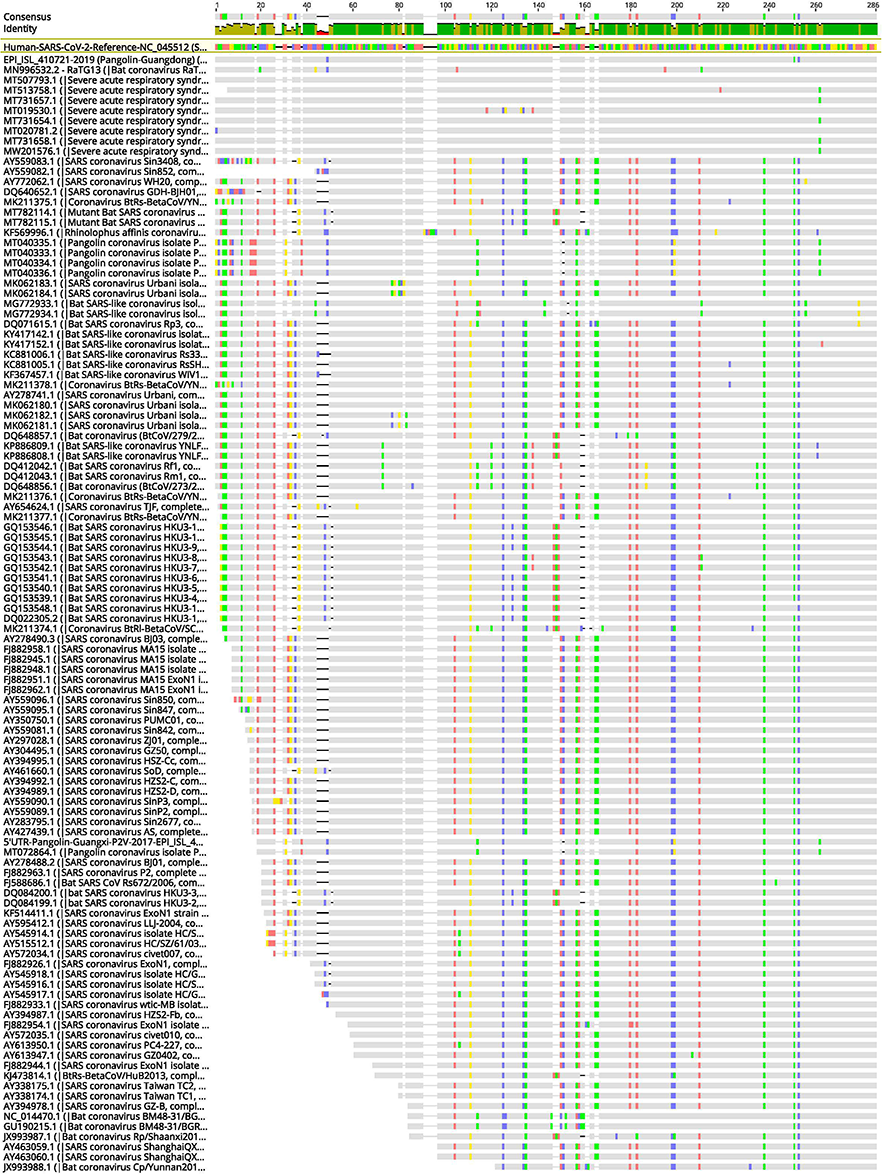
Section of a multiple alignment of the 5’UTR of 505 coronaviruses from diverse hosts.

**Supplementary Table 1 –** List of accession numbers of all viral genomes used in this study.

**Supplementary Table 2 –** D-values of dinucleotide and trinucleotide motifs used as an input for the first PCA.

**Supplementary Table 3 –** D-values of dinucleotide and CpG trinucleotide motifs of all coronaviruses used in this study.

**Supplementary Table 4 –** Zap (ZC3HAV1) mRNA expression level (TPM) in the common reservoir of selected viruses.

**Supplementary Table 5 –** Abundance of C(n7)G(n)CG motif in coronaviruses used in this study.

**Supplementary Table 6 –** CpG D-values of HIV-1 isolate sequences.

## Data Availability

The data underlying this article are available at the online supplementary material.

## Contributions

AA, DE, and HAR conceived the project. HAR and AA conducted all analysis. HAR generated Figures 1 and 2. AA generated all other figures and tables with the supervision of DE and HAR. All authors contributed to the discussions. AA and DE drafted the manuscript, which was reviewed by all authors. All authors read and approved the final manuscript.

